# Whom Do We Prefer to Learn From in Observational Reinforcement Learning?

**DOI:** 10.1101/2025.05.19.654209

**Authors:** Gota Morishita, Carsten Murawski, Nitin Yadav, Shinsuke Suzuki

**Affiliations:** Centre for Brain, Mind and Markets, The University of Melbourne, Melbourne, Victoria, Australia.; Faculty of Social Data Science, Hitotsubashi University, Tokyo, Japan.; HIAS Brain Research Center, Hitotsubashi University, Tokyo, Japan.

## Abstract

Learning by observing others’ experiences is a hallmark of human intelligence. While the neurocomputational mechanisms underlying observational learning are well understood, less is known about whom people prefer to learn from in the context of observational learning. One hypothesis posits that learners prefer individuals who exhibit a high degree of decision noise (randomness in action selection), ’free riding’ on the costly exploration of others. An alternative hypothesis suggests that learners prefer individuals with low decision noise, and imitate consistent and reliable behavior. In a preregistered experiment, we found that most participants preferred to learn from low-noise individuals. Furthermore, exploratory analyses revealed that participants who preferred lownoise individuals tended to rely on imitation of others’ actions, whereas those who preferred high-noise individuals relied more on learning from others’ reward outcomes. These findings offer a potential computational account of how learning styles shape partner selection in social learning.

**Author Summary:** In our daily lives, we often learn by watching others. For example, when starting a new job, we might watch an experienced colleague to learn effective strategies, or an inexperienced coworker to avoid common mistakes. While previous studies have examined how people learn by observing others, less is known about how we decide whom to observe. This study explored whether people prefer to learn from individuals who make consistent choices or those who behave more randomly. At first glance, the answer seems obvious: we would naturally prefer to learn from those who make consistent, reliable decisions. However, there can also be value in learning from someone who behaves unpredictably. For example, when searching for a good restaurant, observing an adventurous friend who tries unfamiliar places might help us discover hidden gems. Despite this potential advantage, we found that most participants preferred consistent decision-makers. Further analysis revealed that participants who favored reliable partners tended to imitate their actions, while those who chose more exploratory partners focused more on outcomes. Our findings suggest that personal learning styles shape partner preferences. These insights could help us understand how people choose whom to learn from in everyday settings like classrooms, or workplaces.

## 1 Introduction

Observational learning, the process through which individuals acquire knowledge and skills by observing others, is a critical ability for social animals, including humans, to survive in uncertain environments [1, 2]. For instance, Adelie penguins utilize observational learning to determine whether there are any predators in the sea before entering the water themselves to forage for food [3]. Instead of diving in right away, they watch how the first penguin reacts after diving in the water. If no predator appears, the rest follow, using this observation to judge whether the environment seems relatively safe. Similarly, humans often rely on observational learning in decision-making processes. For example, when choosing a restaurant, people often read online reviews or consider friends’ recommendations using others’ experiences to identify high-quality restaurants while minimizing the risk of choosing a poor dining experience [4].

Given the ubiquity of observational learning, numerous studies across disciplines have investigated its computational and neural mechanisms [5, 6, 7, 8, 9, 10, 11, 12, 13, 14, 15, 16]. Research has shown that individuals use both reward outcomes and actions of others to guide their own decisions [8, 9, 11]. Specifically, when making choices, people not only rely on action values learned through observing the reward outcomes associated with others’ actions (learning from reward outcomes of others), but they also exhibit a separate bias toward actions frequently chosen by others (learning from actions of others). However, most of these studies predetermined the partners that participants observed, offering limited insight into how learners choose whom to learn from—a crucial aspect in real-world settings, where individuals often have the autonomy to decide whom to learn from. For instance, in workplace settings, individuals may selectively observe experienced colleagues to identify actions that lead to good results, or less experienced peers to recognize and avoid ones that lead to bad results. Nonetheless, little is known about individuals’ preferences for selecting observational partners.

To fill this gap, we examined individuals’ preferences for observational partners, specifically focusing on their preferences regarding a partner’s level of decision noise (i.e., randomness in action selection). From a computational perspective, decision noise modulates the degree of random exploration [17]: higher decision noise leads to greater exploration, whereas lower decision noise corresponds to reduced exploration. In sequential value-based decision-making, individuals face a fundamental trade-off between exploring unfamiliar options to gain information and exploiting familiar options to maximize immediate rewards [18]. Thus, observing partners with different levels of decision noise may influence how individuals learn and adjust their own decision-making, as partners with higher or lower decision noise provide varying amounts of information about available options. Moreover, individual differences in learning styles may lead to differential selection of observational partners based on their decisionnoise characteristics.

We hypothesized that individuals would prefer high-noise partners (i.e., partners who exhibit a high degree of exploration). Learning from such partners may provide advantages by allowing observers to gain information about a wider range of unfamiliar options without directly incurring risk. Economic theory supports this notion, demonstrating that observing highly explorative partners can yield greater informational benefits, ultimately leading to higher total rewards [19, 20].

This hypothesis rests on the assumption that individuals primarily adopt a reward learning style, that is, they infer action values by observing a partner’s reward outcomes. However, if individuals instead rely on an imitative learning style, in which learning is based on copying others’ actions rather than estimating action values using observed outcomes, their partner preferences may differ. In this case, observers may prefer low-noise partners, as such individuals tend to exhibit more consistent and deterministic decision patterns, making them more reliable models for imitation. Indeed, research on social learning indicates that individuals selectively imitate models perceived as successful or reliable [21]. This perspective suggests an alternative hypothesis: when individuals primarily rely on imitation, individuals may show a preference for low-noise partners.

To distinguish between these competing hypotheses, we designed a novel behavioral experiment in which participants chose between two potential partners categorized as either high-noise or low-noise. Participants subsequently engaged in an observational learning task with their selected partner. To ensure reproducible results, we employed a two-step methodological approach. First, we conducted a pilot experiment and performed a power analysis to determine the necessary sample size for the main experiment. Then, we preregistered our analysis plans and conducted the main experiment, collecting data specifically designed to robustly test our hypotheses.

We found that the majority of participants exhibited a preference for selecting a low-noise partner both in the main and pilot studies. To further examine individual differences in partner selection and learning styles, we conducted model-free and model-based exploratory analyses to identify distinct learning styles employed by participants. These analyses revealed that individuals who preferred low-noise partners demonstrated a greater reliance on imitative learning. These findings offer a computational perspective on the mechanisms underlying partner selection in social learning contexts, shedding light on how individuals determine whom to observe for informed decision-making.

## 2 Results

### 2.1 Overview of experimental design

The behavioral experiment comprised four blocks (see Methods for details). In each block, participants completed the Passive Observation task twice, and the Partner Selection and Observational Learning tasks once each (Fig. 1a–d). Prior to these four blocks, participants performed the Individual Learning task as a practice block, allowing them to become familiar with the basic task structure (see Methods for details).

**Figure 1:**
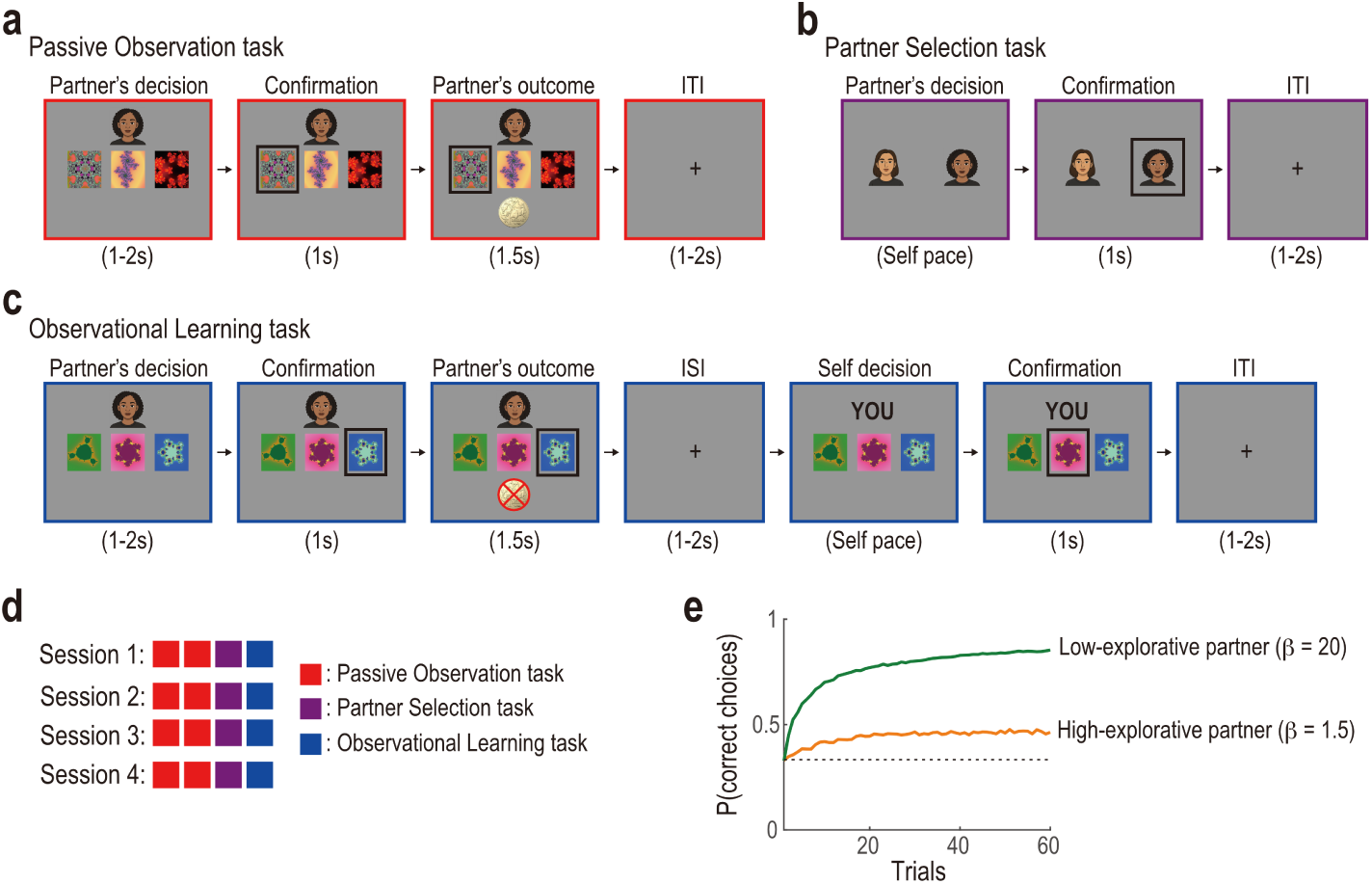
(a) Timeline of each trial in the Passive Observation task. Participants observed a potential partner choose between three options and receive a reward or not. (b) Partner Selection task. Participants chose between two potential partners they had observed in the Passive Observation task as the partner with whom they would perform the Observational Learning task. (c) Timeline of each trial in the Observational Learning task. Participants observed the partner’s choice and outcome, and then made their own choice between the same set of options, without observing their own outcome. (d) Timeline of the entire experiment. In each block, participants completed two Passive Observation tasks, observed two potential partners with differing levels of decision noise, and then performed a Partner Selection task. Finally, they completed an Observational Learning task with the selected partner. The order was randomized across participants. (e) Proportions of correct choices (i.e., choosing the option with the highest reward probability) generated by a reinforcement learning algorithm with different degrees of decision-noise (i.e., inverse temperature *β*). The green line represents the low-noise partner’s learning curve (*β* = 20.0), while the orange line represents the high-noise partner’s curve (*β* = 1.5).

In each of the two Passive Observation tasks, participants observed one potential partner performing a three-armed bandit task (i.e., repeatedly choosing one of the three options and receiving a reward based on the probability associated with the chosen option; Fig. 1a) [22, 23, 24]. The potential partner differed between the first and second Passive Observation tasks: one demonstrated a high degree of decision noise, while the other exhibited a low degree of decision noise (Fig. 1e). The presentation order of the two potential partners and their association with face images were counterbalanced across participants. In the Partner Selection task, participants selected one of the two potential partners to learn from in the subsequent Observational Learning task (Fig. 1b). During the Observational Learning task, participants performed a three-armed bandit task side by side with their chosen partner (Fig. 1c). On each trial, participants observed their partner’s action (i.e., selection of an option) and its outcome (i.e., rewarded or not) before making their own choice. Notably, the outcomes of the participants’ own choices were not displayed, thereby preventing learning from direct reward experiences.

### 2.2 Basic behavior

We first confirmed that participants learned to select the most rewarding option, verifying their comprehension of the task structure. The proportion of correct choices (i.e., choosing the option with the highest reward probability) significantly exceeded chance levels in the practice Individual and Observational Learning tasks, as well as in the main Observational Learning task (Fig. S1: average proportion over 60 trials = 0.73 ± 0.03 (Mean and SEM across participants), *t*(55) = 12.99*, p <* 0.001 for the practice Individual Learning task; proportion = 0.76 ± 0.03*, t*(55) = 13.78*, p <* 0.001 for the practice Observational Learning task; and proportion = 0.71 ± 0.02*, t*(55) = 15.59*, p <* 0.001 for the main Observational Learning task).

### 2.3 Partner selection for observational learning

We next examined the main research question of whom individuals prefer to learn from in the context of observational learning. We found that the majority of participants preferred the low-noise partner over the high-noise partner (Fig. 2). Pilot data (*N* = 20) showed that in the Partner Selection task, 13 out of 20 participants were more likely to select the low-noise partner (i.e., the proportion of selecting the high-noise partner across the four blocks was less than 0.5: see the green bars in Fig. 2a), 5 participants selected the high-noise partner (i.e., the proportion was greater than 0.5: see the orange bars in Fig. 2a), and 2 participants exhibited no preference (i.e., the proportion was equal to 0.5: see the white bar in Fig. 2a).

**Figure 2:**
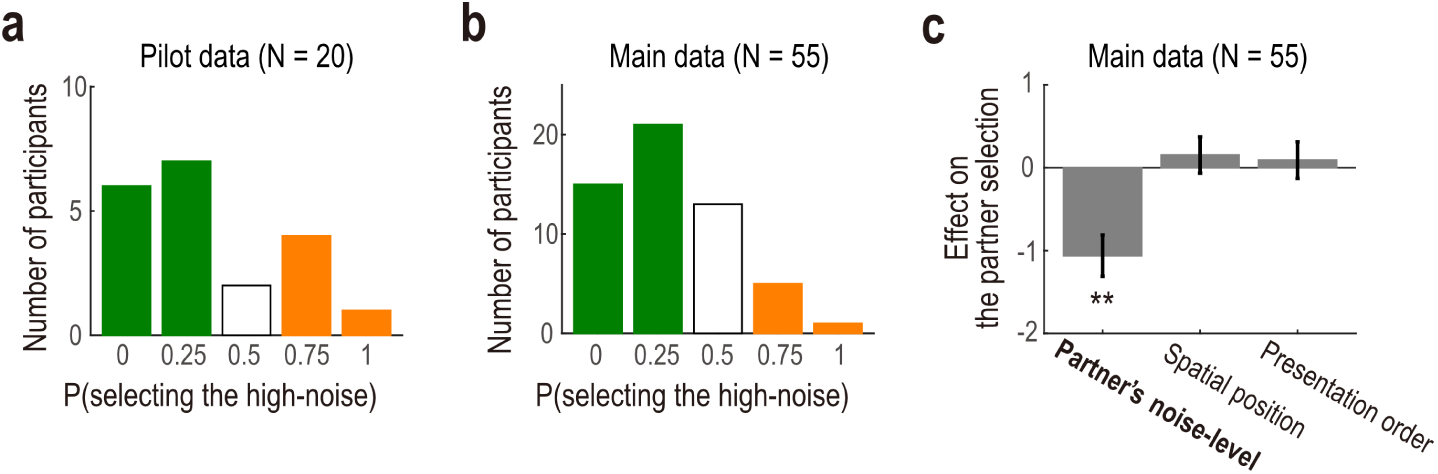
Partner preference results. (a) Histogram of the proportions of selecting the high-noise partner for the pilot data (*N* = 20). (b) Histogram of the proportions of selecting the high-noise partner for the main data (*N* = 55). In both (a) and (b), bar colors represent levels of preference: green bars indicate a low preference for selecting the high-noise partner, the white bar represents neutral preference (0.5), and orange bars indicate a high preference for selecting the high-noise partner. (c) Effects on partner selection in the main study (Mean ± SEM, *N* = 55). The leftmost bar shows a significant negative effect of the partner’s exploration level on selection probability (*p <* 0.001), indicating that participants were less likely to choose the high-noise partner. The other bars show non-significant effects of spatial position and presentation order. The means and SEMs were estimated with a generalized linear mixed model (GLMM). p-values were obtained using two-tailed t-tests.

To rigorously evaluate the findings of the pilot study, we conducted a preregistered main experiment. The sample size (*N* = 55) was determined through a formal power analysis (see Methods for details). Consistent with the pilot data, participants in the main experiment demonstrated a preference for the low-noise partner in the Partner Selection task (Fig. 2bc). That is, most participants were more likely to select the low-noise partner to learn from for the subsequent Observational Learning task (proportion of selecting the high-noise partner over the four blocks was less than 0.5; Fig. 2b). A mixed-effects logistic regression analysis revealed a significant negative effect of the partner’s decision-noise level on partner selection (*b* = −1.06 ± 0.25*, t*(55) = −4.27*, p <* 0.001; Fig. 2c), even after controlling for potentially confounding effects of the spatial position of the partner (i.e., whether the potential partner was displayed on the left or right side of the screen in the Partner Selection task; see Fig. 1b) and the presentation order (i.e., whether the potential partner was presented in the first or second Passive Observation task). These results collectively suggest that most participants preferred to learn from the low-noise partner for observational learning.

### 2.4 Individual difference in the partner selection and style of observational learning

The preregistered experiment above revealed that the majority of participants preferred learning from the low-noise partner, while others preferred the highnoise partner. What mechanism underlies the individual difference in partner selection for observational learning? We reasoned that these differences reflect participants’ learning styles. Specifically, participants who preferred the lownoise partner may have relied more on learning from the partner’s actions (i.e., imitation), as low-noise partners consistently made good choices, making the partners more consistent and reliable for imitation. In contrast, participants who preferred the high-noise partner may have relied more on learning from the partner’s reward outcomes, since high-noise partners explored a broader range of options, enabling observers to learn about the rewards associated with different choices and discover better options themselves. To test this reasoning, we conducted exploratory analyses combining the pilot and main experiment data together. In the exploratory analyses, effect sizes (e.g., standardized regression coefficients and Cohen’s d) are reported instead of p-values, as p-values are more appropriate for statistical judgments based on a priori hypotheses [25, 26].

Using a generalized linear mixed model (GLMM), we assessed the extent to which each participant relied on learning from the partner’s action and reward, as well as how this learning style was related to individual differences in partner selection. The GLMM quantified that, in the Observational Learning task, how much participants’ trial-by-trial behavior was influenced by the partner’s past action and reward. In addition to the main effects of the partner’s past action and reward, the GLMM included the interaction terms with the individual difference in partner selection (i.e., the proportion of blocks in which the participant selected the high-noise partner in the Partner Selection task). Crucially, the interaction terms allowed us to quantify how learning styles (i.e., the effects of the partner’s past action and reward) were associated with partner preferences across participants.

The GLMM revealed that, on average, participants in the Observational Learning task learned from both the partner’s past action and reward. The partner’s past action and reward had positive effects on participants’ behavior (*b* = 0.83 ± 0.06 for the action effect; and *b* = 1.63 ± 0.11 for the reward effect; Fig. 3a), consistent with prior research on observational learning [8, 9, 15]. Moreover, the effects of the partner’s past action on participants’ behavior in the Observational Learning task were modulated by individual differences in partner selection, as indicated by the interaction effect (Fig. 3a). That is, the effect of the partner’s past action was negatively modulated by the participants’ proportion of selecting the high-noise partner (*b* = −1.69 ± 0.15). The interaction effect supports our reasoning that participants who selected the low-noise partner relied more on learning from the partner’s past action compared with those who selected the high-noise partner.

**Figure 3:**
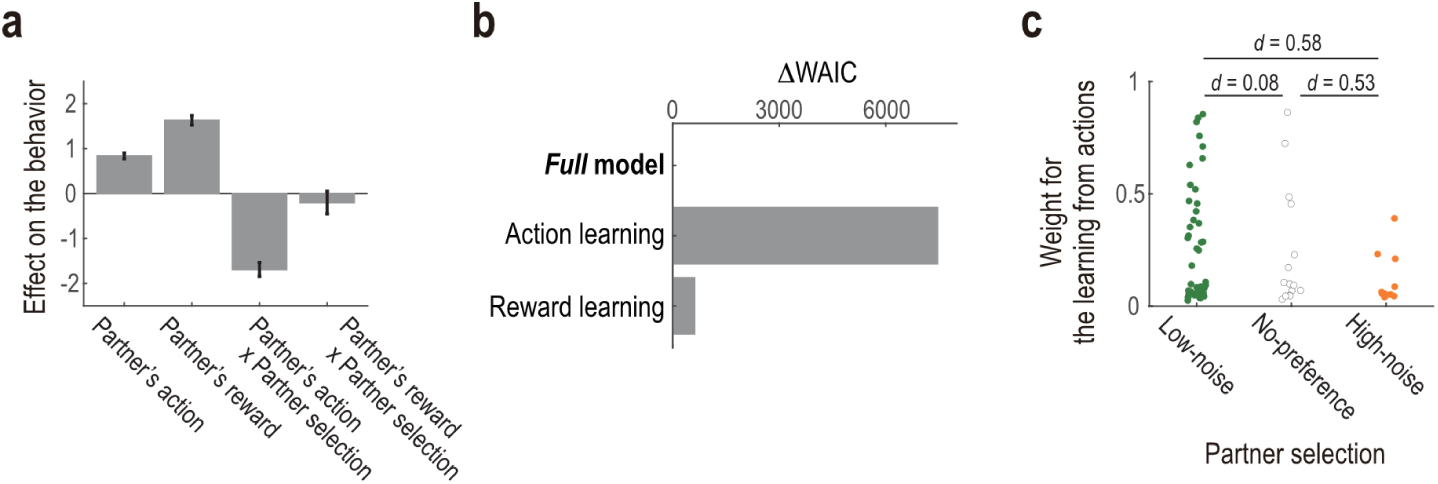
Exploratory analyses of individual learning styles. (a) Effects of partner-related variables on participants’ behavior (Mean ± SEM, N = 74). The first and second bars represent the main effects of the partner’s action and reward, respectively. The third bar shows the interaction between partner’s action and partner selection (defined as the proportion of blocks in which the high-noise partner was chosen), indicating that imitation behavior varied with partner preference. The fourth bar reflects the interaction between partner’s reward and partner selection. The means and SEMs were estimated with a generalized linear mixed effects model (GLMM). We did not report p-values as the regression analysis was explanatory. (b) Model comparison using Widely-applicable Akaike Information Criteria (WAIC). The full model, which includes both action and reward learning styles, provides the best fit to the data.(c) Estimated individual weights for learning from partner actions extracted from the Full model, grouped by partner preference. Cohen’s d values indicate effect sizes between groups.

This reasoning is further supported by computational modeling. We compared three computational models of observational learning based on their goodness of fit to participants’ actual choice data in the Observational Learning task. We then compared parameter estimates in the best-fitted model between participants who preferred to learn from the low-noise partner and those who preferred the high-noise partner.

Inspired by previous studies on observational learning [15, 16, 27], our full model assumes that an agent combines two learning systems with a certain weight: learning from the partner’s actions (i.e., imitation) and learning from the partner’s rewards. Note that the main parameter of interest in the full model is the weight of the two learning systems. We also tested for alternative partial models that include only one of the two learning systems (i.e., the action learning model and reward learning model).

Those models were fitted using a hierarchical modeling approach [28, 29], and their goodness of fit was assessed based on the Widely-applicable Akaike Information Criterion (WAIC) [30]. The model-fitting procedure was validated through model-recovery and parameter-recovery analyses on synthesized data [31]. The model-recovery analysis confirmed the identifiability of the competing models: simulation data generated by each model were better captured by the corresponding model than by the other models, as evidenced by the confusion matrix being close to the identity matrix [31] (Fig. S2a). The parameter-recovery analysis demonstrated that estimated values of the model parameters aligned well with the true generative values used in the simulation (Fig. S2b). These simulation results support the validity of our model-fitting procedure.

The model comparison indicated that the full model provided a better fit compared to the other competing partial models (Fig. 3b). This finding suggests that participants employed both learning from the partner’s actions and rewards for decision-making in the Observational Learning task (see Fig. S3 for posterior densities of the group-level mean parameters), consistent with the results of the GLMM analysis (Fig. 3a) and prior studies [8, 9, 15]. Exploring the individual difference in the parameter estimate of interest in the best-fitting full model, we then found that the weight assigned to learning from the partner’s actions (relative to that from the rewards) was greater in participants who preferred the low-noise partner compared to those who preferred the high-noise partner (Cohen’s d = 0.58; see Fig. S4 for comparison of other parameters; Fig. 3c). These results from the model-based analysis were consistent with our reasoning that participants who preferred the low-noise partner more, learned from the partner’s actions (compared with those who preferred the high-noise partner) and that those who preferred the high-noise partner more, learned from the partner’s rewards.

## 3 Discussion

Observational learning is a fundamental cognitive ability that enables individuals to acquire knowledge and skills by observing others. Given its significance, numerous studies across various disciplines have examined the mechanisms underlying observational learning. However, few studies have directly addressed the critical question of whom individuals prefer to learn from. To address this question, we developed a novel behavioral paradigm where participants selected between two potential learning partners who differed in their level of decision noise for observational learning. To ensure the reproducibility of our findings, we preregistered our analysis approach and based our sample size on prior pilot data. The preregistered analysis revealed that participants, on average, preferred to learn from the partner with lower decision noise. Building on this, we examined whether and to what extent individuals relied on their partner’s actions (i.e. imitation) or reward outcomes during the observational task. We found that participants relied on both action-based and reward-based learning styles, consistent with previous studies [8, 9, 15]. More importantly, participants who preferred low-noise partners were more likely to rely on imitation, suggesting that personal learning styles may systematically shape preferences for social learning partners.

The finding that participants preferred a low-noise partner might initially seem intuitive and straightforward. However, from a learning and information acquisition perspective, lower decision noise, meaning less exploration, can hinder effective learning. In many learning contexts, especially those involving uncertainty, exploration is essential for identifying which actions yield the highest rewards. Without exploration, learners risk overlooking better alternatives. However, exploration also entails risks and, more importantly, can lead to missed opportunities for exploiting the option currently believed to be the best. A fundamental advantage of observational learning is that individuals can learn about the consequences of actions without being exposed to the risks associated with exploration or sacrificing exploitation opportunities. From this normative standpoint, observing a high-noise partner should be beneficial, as higher decision noise indicates greater exploration, offering more opportunities to learn about unfamiliar options without personally incurring exploration risks. Indeed, formal economic models of strategic learning support this view, demonstrating that individuals can optimize their own reward outcomes by minimizing personal exploration when observing a partner who exhibits a higher degree of exploration [19, 20]. These theoretical predictions raise an important question: why did participants in our study prefer a low-noise partner?

One possible explanation is that, while exploration is necessary to solve the three-armed bandit task used in our experiment, participants may have perceived that extensive exploration was not essential for optimal performance. If the task structure did not strongly incentivize exploration, participants may have been less motivated to select a high-noise partner. Future research could further investigate this possibility by modifying the task to increase the necessity of exploration. For instance, associations between choices and reward probabilities could be swapped midway through the task, or the reward probabilities systematically altered over time. These manipulations would potentially make a high-noise partner more valuable.

Another alternative explanation is that participants may have favored the low-noise partner because their behavior more closely matched their own decisionmaking style. In our data, participants’ learning curves during the individual task looked very similar to those of the low-noise partner (Fig. S1a vs. Fig. 1e), suggesting a match in behavior. This is in line with recent research showing that learners benefit more when learning from demonstrators whose actions are predictable [14], suggesting that perceived behavioral similarity may enhance the effectiveness of social learning. While the present study did not directly measure participants’ perceptions of how closely their partners’ decisions aligned with their own, future studies could include such measures to better understand how perceived similarity affects partner choices.

Another interpretation is that participants may have simply chosen the partner who performed better. In our experiment, the low-noise partner earned more reward than the high-noise partner (Fig. 1e), although this is not necessarily always the case, as a low-noise partner might ultimately settle on a suboptimal choice due to insufficient exploration. Prior research has shown that individuals tend to learn more quickly and effectively when learning from someone who is skilled or successful, as opposed to someone who performs poorly or inconsistently [32, 33, 34, 35]. In our design, we manipulated the level of decision noise (i.e., exploration level), but exploration and performance were inherently linked—lower decision noise led to better performance—making it difficult to determine whether participants’ preferences were driven by the partner’s exploration style or by their observed success. To disentangle these factors, future research should aim to decouple exploration from performance. For example, the task could be structured such that partners with varying levels of exploration achieve comparable performances. This approach would allow researchers to isolate the effect of exploration itself on partner selection, eliminating the confounding influence of performance differences.

Another possible account is that participants may have been motivated to minimize cognitive effort. Observing a high-noise partner likely imposes greater demands on working memory, as it requires tracking a larger variety of actions and their associated outcomes. Prior work has shown that models incorporating memory constraints, such as forgetting-based Q-learning algorithms, often provide a better fit to human behavior in sequential decision-making tasks [36, 37, 38, 39, 40]. This suggests that individuals may struggle to accurately retain and update information when the number of distinct observations increases. From this perspective, a low-noise partner may be preferred simply because their behavior is easier to follow and integrate.

Some individuals may adopt simpler social learning styles, such as imitative learning, which place fewer cognitive demands on memory and computation compared to reward learning styles [41, 42]. If so, individuals relying on imitation should be especially likely to favor low-noise partners, whose behavior is not only more predictable but also more likely to choose the best-known option, making them both easier and more advantageous to imitate. This interpretation is supported by our finding that participants who preferred the low-noise partner were more likely to learn from the partner’s actions.

A similar but distinct account involves social conformity or peer influence, which can lead individuals to imitate others’ behavior regardless of whether it is the most informative strategy [43, 44, 45, 46, 47]. In our task, participants were able to observe the reward outcomes of the partner’s actions, so it is reasonable to assume that a reward learning style may have guided their choices. Even so, if participants instead relied on imitation, selecting a low-noise partner would likely have led to higher rewards, as that partner consistently chose the most rewarding option. This interpretation aligns with previous research showing that people tend to selectively imitate behaviors that are performed consistently and reliably by others [32, 33, 34, 35].

To further probe this possibility that people who preferred a low-noise partner actually tended to imitate the partner’s actions, we employed computational modeling analysis as well as model-free regression analysis. The results indicate that participants indeed differed in their learning styles based on partner preferences. Participants who preferred the low-noise partner had a higher weight on the Action Learning component (see Methods for details), relying more on learning from their partner’s actions rather than learning from their partner’s reward outcomes, compared to those who preferred the high-noise partner (Fig. 3c). However, this analysis cannot conclusively determine whether this preference was driven by the participants’ inherent learning style or by their partner’s specific behaviors. To address this question, future research should include an experiment where participants are randomly paired with partners exhibiting varying degrees of decision noise. Such a design would help clarify whether learning styles are inherently linked to partner preferences or primarily driven by observed partner behavior.

An intriguing direction for future research might be exploring how social and psychological traits may impact partner selection in value-based decision making. Specifically, attributes such as perceived trustworthiness or ideological alignment could meaningfully shape preferences during observational learning. Prior research across diverse disciplines has highlighted the importance of such factors in shaping partner selection decisions. For example, studies in mate preference have consistently demonstrated the significance of traits such as trustworthiness, attractiveness, and status in romantic partner selection [48, 49]. In songbird studies, juvenile male zebra finches exposed to two adult males exhibited a preference for learning songs from tutors who displayed more aggressive behavior toward them [50]. Although not focused explicitly on preference, research in observational learning has shown that individuals are more likely to learn from others who share their political views [12]. Despite the extensive literature on partner preference in other domains, limited attention has been devoted to understanding the role of these social and relational attributes in decision-making contexts. Investigating how attributes such as trustworthiness or political alignment affect partner selection in observational learning could yield critical insights into the mechanisms of social learning and decisionmaking.

Another promising direction for future research involves investigating how preferences for learning partners evolve over time through repeated interactions. In the current experimental design, participants made a one-time choice between two potential partners and did not return to those options in later blocks. However, in real-world social learning scenarios, individuals often have repeated opportunities to choose whom to learn from, allowing for the gradual development of preference for a partner over time. For instance, in workplace environments, employees may initially consult multiple colleagues when tackling unfamiliar tasks. Over time, they tend to rely more on those who consistently provide accurate or helpful advice, leading to the emergence of trusted collaborators. This process reflects a dynamic updating of partner value based on repeated interactions and feedback. Investigating how such long-term relationships form and influence partner preference could provide valuable insights into the mechanisms of social learning.

In conclusion, our study investigated whom individuals prefer to learn from in observational learning contexts and found a clear preference for low-noise partners, contrary to theoretical predictions emphasizing the benefits of observing high-noise partners. Our computational modeling analyses suggest that this preference might be driven by tendency to imitate. We believe these findings offer valuable insights into the computational mechanisms underlying partner selection, contributing to our understanding of how social learning networks might be constructed and how collective behaviors emerge in social groups.

## 4 Methods

### 4.1 Ethics statement

The experimental protocol was approved by the University of Melbourne Human Research Ethics Committee (Ethics ID 26997). All participants provided written informed consent before the experimental session commenced.

### 4.2 Preregistration

The experimental plan was preregistered on the Open Science Framework (OSF) and can be accessed at https://osf.io/g6etf.

### 4.3 Participants

Participants were recruited through advertisements on the University of Melbourne’s student portal. Eligible participants were aged 18 to 35 years. Each participant received a A$10 show-up fee and additional performance-based compensation. Twenty participants took part in the pilot experiment (15 females, 4 males, and 1 unidentified; age range, 18-28 years; mean age ± SD, 22.5 ± 2.86), and 55 were recruited for the main experiment (35 females, and 20 males; age range, 18-29 years; mean age ± SD, 22.98 ± 3.25). In the pilot study, one participant completed only two of the four blocks; this individual was included in the power analysis and preliminary analyses conducted prior to the main experiment but excluded from subsequent analyses. Participants received instructions via slides and completed comprehension quizzes to ensure they correctly understood the experimental task. Every participant answered all the questions correctly.

Participants were told their partners were previous human participants in both the Passive Observation tasks and the Observational Learning tasks, but in reality, the partners were simulated agents using a Q-learning algorithm (See ”Observational Learning task” and ”Passive Observation task” subsections below for details). After the experiment, participants were debriefed about the deception and could withdraw their data if desired. Additionally, we conducted brief interviews to assess whether participants had suspected that the partner choices were not generated by real individuals. No participants expressed strong doubts regarding the authenticity of their partners.

### 4.4 Experimental design

We conducted both a pilot behavioral experiment and a preregistered confirmatory behavioral experiment, in which participants engaged in observational learning alongside partners they selected themselves (Fig. 2). The experimental protocol consisted of two preparatory blocks followed by four main task blocks.

In the first preparatory block, participants independently performed a threearmed bandit task (see the “Individual Learning task” subsection below). In the second preparatory block, they participated in an observational learning task (see the “Observational Learning task” subsection below) with a pre-determined partner. During each of the main blocks, participants initially observed two potential partners executing the individual learning task (see the “Passive Observation task” subsection below). Following the observation, participants selected one of the observed individuals as their partner for the subsequent Observational Learning task (see the “Partner Selection task” subsection below). Finally, participants engaged in the Observational Learning task with their chosen partner.

#### 4.4.1 Individual Learning task

Participants performed a three-armed bandit task, where they repeatedly chose between three stimuli (fractal images) for 60 trials. Each stimulus was associated with a hidden reward probability of 0.25, 0.5, or 0.75, respectively. The associations between stimuli and reward probabilities were randomized across participants. Participants were explicitly informed that the set of stimuli, their positions on the screen, and the associated reward probabilities would remain constant throughout the block.

#### 4.4.2 Observational Learning task

Participants repeatedly made choices between three stimuli, not based on their reward outcomes, but by observing their partner’s choices and outcomes (Fig. 1c). Similar to the Individual Learning task, the three stimuli were paired with hidden reward probabilities of 0.25, 0.5, and 0.75, respectively. These associations remained constant throughout the block but were randomized across participants. It is important to note that participants were informed that both they and their partner were faced with the same stimulus-reward associations. In each of the 60 trials, participants first observed their partner’s choice and its outcome. The chosen stimulus was highlighted with a black frame for 1 second, followed by the display of the outcome for 1.5 seconds. Afterward, a fixation cross appeared for a variable duration ranging from 1 to 2 seconds. Subsequently, participants made their own choice. The chosen stimulus was also highlighted with a black frame for 1 second. However, unlike their partner’s choice, the outcome of the participant’s choice was not revealed.

Crucially, in the preparatory block, participants observed a pre-determined partner. They were informed that this partner was another individual who had participated in the individual learning task on a previous day. However, the ’partner’ was a simulated agent operating on a standard Q-learning algorithm with a learning rate set to 0.3 and an inverse temperature parameter of 7.0. In contrast, during the main blocks, participants selected their partners (see ”Passive Observation task” and ”Partner Selection task” subsections below).

#### 4.4.3 Passive Observation task

Participants observed two potential partners, each exhibiting different levels of decision noise, which were unknown to participants. The partner with a higher level of decision noise was simulated using a standard Q-learning algorithm with an inverse temperature parameter of 1.5 (Fig. 1e). In contrast, the partner with lower decision noise was simulated with an inverse temperature parameter of 20.0. The learning rate was set to 0.3 for both simulations (Fig. 2e).

To help participants differentiate between the two potential partners, distinct face images were used (all of the same sex, with a multi-racial face image dataset employed to minimize potential sex and racial biases) [51] ^1^. Each participant first observed one potential partner completing 30 trials of the individual learning task, followed by the observation of the other partner. The presentation order of the partners was randomized across the four main blocks, while their corresponding face images were randomized across participants.

#### 4.4.4 Partner Selection task

For the subsequent Observational Learning task, participants chose between two potential partners: a high-noise partner and a low-noise partner, both of whom they had observed in the preceding Passive Observation task. In this task, the two facial images representing the potential partners were displayed side by side in a horizontal layout. The associations between the presentation positions of these images and the exploration levels of the potential partners were counterbalanced across blocks. Participants made their selection by clicking on the face image of the partner they preferred.

#### 4.4.5 Software

We coded the experimental tasks using Node.js (https://nodejs.org/en) in MacOS.

### 4.5 Statistical analysis

#### 4.5.1 Regression analyses

To address the main question of whom participants preferred to learn from during observational learning, we conducted a GLMM analysis—specifically, a mixed-effects logistic regression—using the glmer function from the lme4 package (version 1.1-35) in R [52, 53]. As this analysis was preregistered, we included only data from the main experiment (*N* = 55).

The dependent variable *Y* was coded as 1 if a potential partner displayed on the left-hand side in the Partner Selection task was chosen, or 0 if a potential partner displayed on the right-hand side was chosen. Two independent variables were defined in this regression. The first independent variable was coded as 1 if a potential partner with a higher level of decision noise is presented on the left-hand side, and as −1 otherwise. The coefficient of this variable represents the effect of the partner’s decision-noise level on partner selection. The second independent variable controlled for the presentation order bias. It was coded as 1 if the potential partner displayed on the left-hand side was first presented in the preceding Passive Observation task, and −1 otherwise. The intercept controlled the effect of the spatial position of the partner (see Fig. 1b).

The GLMM was fit to the main data treating a participant as a random factor and allowing for random intercepts and slopes for each independent variable. Point estimates of coefficients and their standard errors were obtained for each fixed effect in the model. Then, two-tailed t-tests were performed to examine whether each coefficient differed statistically significantly from zero.

For the regression analysis without the presentation order and spatial bias controls, a bootstrap-based power analysis [54] using pilot data indicated that a sample size of 55 would achieve a power of 0.8 at a 0.05 significance level (two-tailed).

We investigated the individual learning style, that is, how the partner’s actions and rewards influenced the participant’s current choice within each trial. Note that because this is an exploratory analysis, we combined the pilot data with the main data (*N* = 74). Following previous studies using a three-armed bandit task [22, 23, 24], we ran three separate GLMMs, one for each option (*X*, *Y*, and *Z*). Each GLMM estimated the probability that participants chose option *X* (or *Y*, or *Z*) on each trial. Each model took the following form in Wilkinson notation (using option *X* as an example):

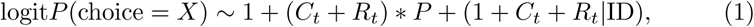

where the dependent variable was coded as 1 if the partner’s choice was *X* on trial *t*, or 0 if the participant chose *Y* or *Z*.

Here, *C_t_*represents the partner’s recent choice, coded as +1 if the partner chose option *X*, −1 if the partner chose any other option. *R_t_* captures the partner’s recent outcome on trial *t* and was coded as +1 if the partner was rewarded for selecting *X*, −1 if rewarded for a different option, or 0 when no reward was given. The continuous variable *P* represents the participant’s preference for choosing high-noise partners, which was defined as a ratio of the number of blocks where a participant chose a high-noise partner. ID is the participant identity.

We repeated this model specification for option *Y* and option *Z*. We derived a mean effect of each predictor across the three models. Specifically, we obtained the variance-weighted average of its coefficients (and corresponding standard errors) across *X*, *Y*, and *Z* models and its variance. In this exploratory regression analysis, we report standardized regression coefficients as effect sizes rather than p-values, as p-values are more appropriate for statistical inferences based on a priori hypotheses [25, 26].

#### 4.5.2 Computational models

To identify a participant’s learning style for observational learning, we considered three computational models. These models were fitted to the choice data in the Observational Learning task and compared based on their goodness of fit. We then examined differences in parameter estimates from the best-fitting model between participants who chose to learn from the low-noise partner and those who opted for the high-noise partner.

##### Full model

This model integrated two learning styles: learning from a partner’s actions and learning from a partner’s rewards. Specifically, it learned the partner’s action tendencies (i.e., which action the partner tends to choose) from their actions and learned action values of available options from their reward outcomes through reinforcement learning. The model assigns two key quantities to each option *X, Y, Z*:

**Action values** (*V_X_, V_Y_, V_Z_*) representing learned action values from observational learning.
**Action tendencies** (*A_X_, A_Y_, A_Z_*) capturing the tendency to imitate a partner’s actions.

Each of these is initialized as follows: *V_X_* = *V_Y_*= *V_Z_* = 1*/*2 and *A_X_* = *A_Y_*= *A_Z_* = 1*/*3.

The model updates these values separately, following the learning rules of its two components: Reward Learning and Action Learning components. The Reward Learning component updates action values based on observed rewards. If a partner chooses option *X* on trial *t*, the update follows:

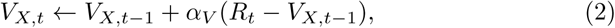

where *R_t_*∈ {0, 1} is the observed reward, and *α_V_* ∈ [0, 1] is the learning rate. The Action Learning component updates action tendencies based on a partner’s choices, reinforcing chosen actions while decreasing the tendency for unchosen ones:

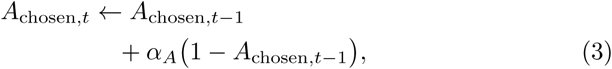

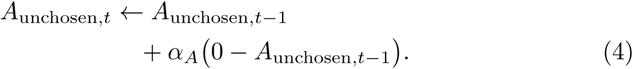

Here, *α_A_*∈ [0, 1] is the learning rate for action tendencies.

To determine which option to choose, the model combines both learned action values and action tendencies using a weighting parameter *w_A_* ∈ [0, 1]:

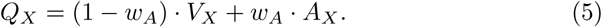

The probabilities of selecting an option were calculated using the softmax rule:

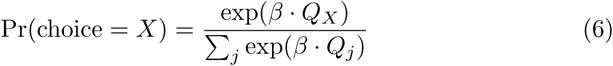

where *β* ≥ 0 is an inverse temperature parameter controlling decision-noise or random exploration (higher *β* leads to more deterministic choices).

##### Action Learning model

This model only learned from a partner’s actions to imitate their choices. It is a partial model in a way that it lacked the Reward Learning component of full model. The action tendency updates were done in the same manner as full model. The choice probabilities were computed in the same manner except this model used the action tendency *A* instead of the combined value *Q*.

##### Reward Learning model

This model only learned from a partner’s rewards. It is also a partial model because it lacked the Action Learning component compared to Full model. The action value updates occured in the same manner as Full model. The choice probabilities were calculated using the softmax rule except this model only used the action value *V* instead of *Q*.

#### 4.5.3 Model fitting

To fit the different computational models to participants’ choice data in the Observational Learning task, we employed a Bayesian hierarchical modeling approach. This was done using Markov chain Monte Carlo (MCMC) sampling implemented in Stan (version 2.35.0) via the Python interface CmdStanPy (version 1.2.1). Four independent chains were run with 1,000 warm-up iterations followed by 1,000 sampling iterations, resulting in a total of 4,000 posterior samples per parameter.

In this hierarchical framework, individual-level parameters are first defined on an unconstrained (real-valued) scale and then transformed to respect their respective domains. For parameters constrained to the unit interval, such as the learning rate for imitation *α_A_*, we define raw parameters *α_A,_*_raw_ on the real-valued line and apply a logistic transformation:

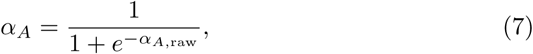

to ensure that *α_A_* ∈ [0, 1]. For strictly positive parameters, such as the inverse temperature *β*, we define a raw parameter *β*_raw_ on the real-value line and map it via an exponential transformation:

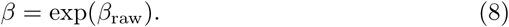

At the group level, the raw parameters are modeled as drawn from normal distributions:

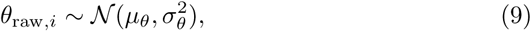

where *θ* stands in for any transformed parameter (e.g., *α_A_*, *α_V_*, or *β*). Hyper-parameters are given weakly informative priors:

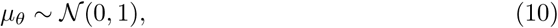

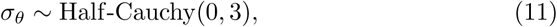

a choice that enforces non-negativity while allowing for substantial variation in individual-level estimates [28].

#### 4.5.4 Model Recovery Analysis

To ensure the robustness and reliability of our model comparison, we performed a model recovery analysis [31]. Simulated choice data were generated using each model under experimental conditions mirroring the study design (74 participants, 60 trials per block, and four blocks). The simulated data assumed that the probability of a partner being low explorative was 0.7, and the probability of a partner being high explorative was 0.3, reflecting the empirical partner choice distribution (Fig. 2ab). Parameter values were drawn from Beta distributions (Beta(1.1, 1.1)) for learning rates and the weight and a Uniform distribution (*U* (0, 30)) for the inverse temperature.

Each simulated dataset was then fit using all three candidate models, and WAIC was calculated to identify the best-fitting model. This procedure was repeated 30 times, allowing us to construct a confusion matrix representing the proportion of times the generating model was correctly identified (Fig S2 a).

#### 4.5.5 Parameter Recovery Analysis

To further assess the validity of our parameter estimates, we conducted a parameter recovery analysis using the Full model, which was identified as the best-fitting model. Simulated choice data were generated using parameter values drawn from the same group-level distributions described above. The full model was then fit to the simulated data using the same MCMC procedure.

We were interested in individual weighting parameters for imitation *w_A_* (see Computational models), so we evaluated individual parameter recovery by computing Pearson correlation coefficients between the true simulated parameters and the corresponding estimated parameters. This process was repeated 40 times to ensure robust results. Strong positive correlations would indicate reliable parameter recovery, supporting the validity of the model’s parameter estimates (Fig. S2b).

## 5 Author Contributions

- **Conceptualization:** Gota Morishita, Shinsuke Suzuki.
- **Data curation:** Gota Morishita, Shinsuke Suzuki.
- **Formal analysis:** Gota Morishita, Shinsuke Suzuki.
- **Funding acquisition:** Carsten Murawski.
- **Investigation:** Gota Morishita.
- **Methodology:** Gota Morishita, Shinsuke Suzuki, Carsten Murawski, Nitin Yadav.
- **Project administration:** Gota Morishita, Shinsuke Suzuki, Carsten Murawski.
- **Resources:** Gota Morishita, Shinsuke Suziki, Carsten Murawski.
- **Software:** Gota Morishita, Shinsuke Suzuki.
- **Supervision:** Shinsuke Suzuki, Carsten Murawski, Nitin Yadav.
- **Validation:** Gota Morishita, Shinsuke Suzuki, Carsten Murawski, Nitin Yadav.
- **Visualization:** Gota Morishita, Shinsuke Suzuki.
- **Writing – original draft:** Gota Morishita, Shinsuke Suzuki.
- **Writing – review & editing:** Gota Morishita, Shinsuke Suzuki, Carsten Murawski, Nitin Yadav.

## 6 Data availability statement

All data, and statistical analysis code will be available on Zenodo at https://doi.org/10.5281/zenodo.15386571.

---

**Figure S1:**
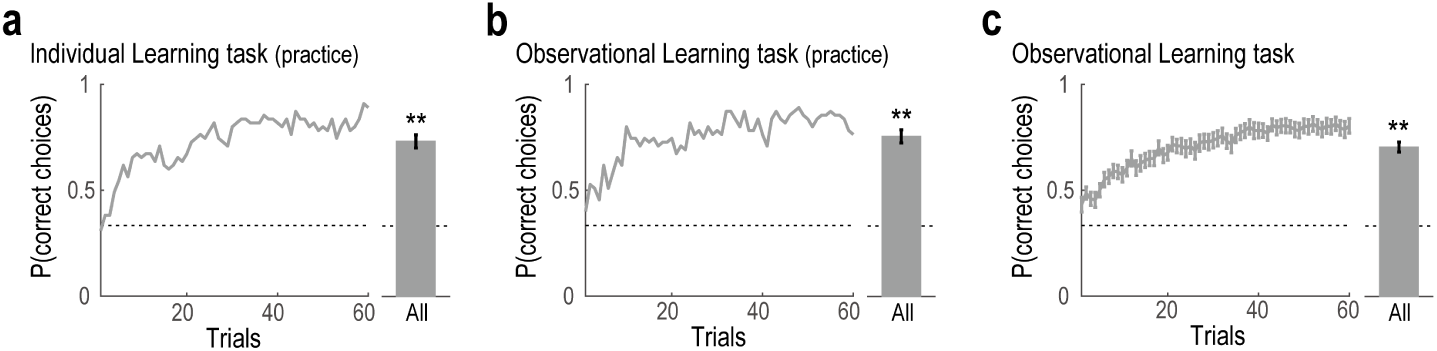
The proportions of correct choices. (a) In Individual Learning task, the average proportion of correct choices increased over the course of trials. (b) In Observational Learning task in the practice block, the average proportion of correct choices increased over the course of trials. (c) In Observational Learning task with the selected partner, the average proportion of correct choices (average across 4 blocks ± SEM) increased over the course of trials.

**Figure S2:**
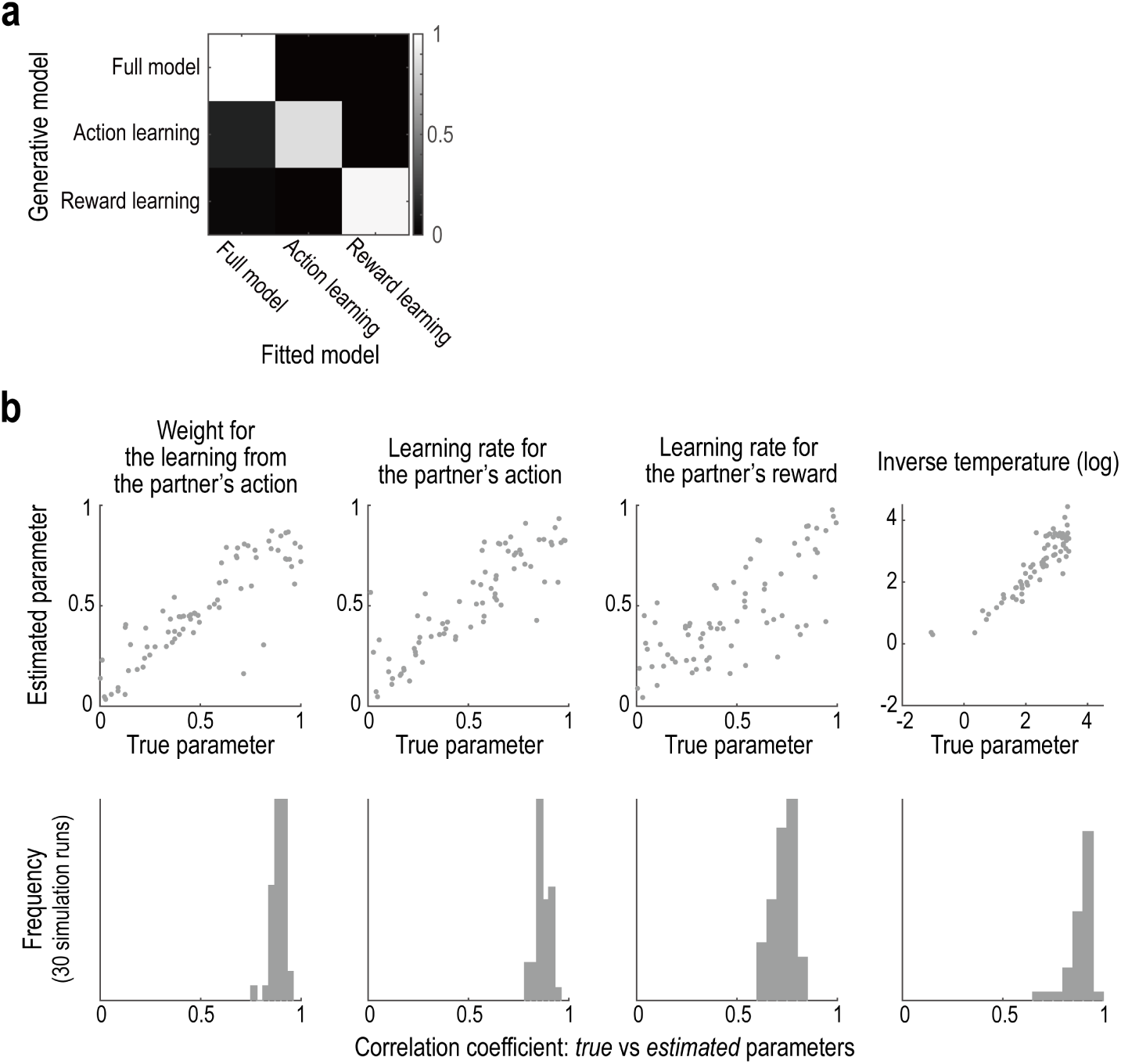
(a) Confusion matrix showing the proportion of times each generative model (rows) was correctly identified by model fitting (columns) based on WAIC. Simulated datasets were generated under conditions matching the empirical design (74 participants, 60 trials per block, 4 blocks) and repeated 30 times. (b) Parameter recovery for the full model. Top: Scatter plots show the correspondence between true (simulated) and estimated values for four parameters: the weight for learning from the partner’s action, learning rates for partner’s action and reward, and the inverse temperature (log scale). Bottom: Histograms of Pearson correlation coefficients between true and estimated parameters across 40 simulations.

**Figure S3:**
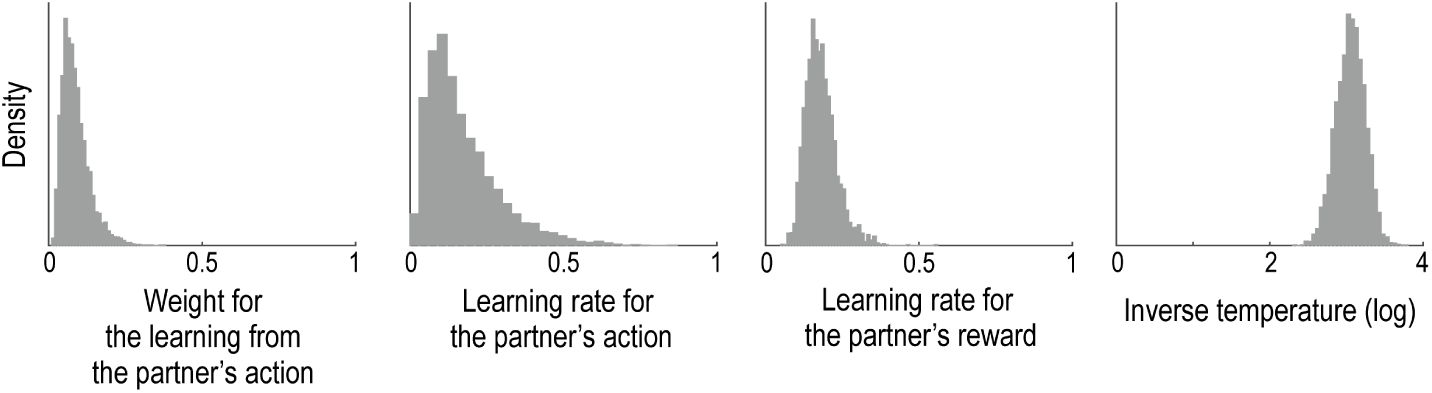
Posterior population-level distributions of parameters in the Full model. Density plots show the group-level posterior distributions for each parameter estimated in the Full model: the weight for learning from the partner’s action, the learning rates for the partner’s action and reward, and the inverse temperature (log scale).

**Figure S4:**
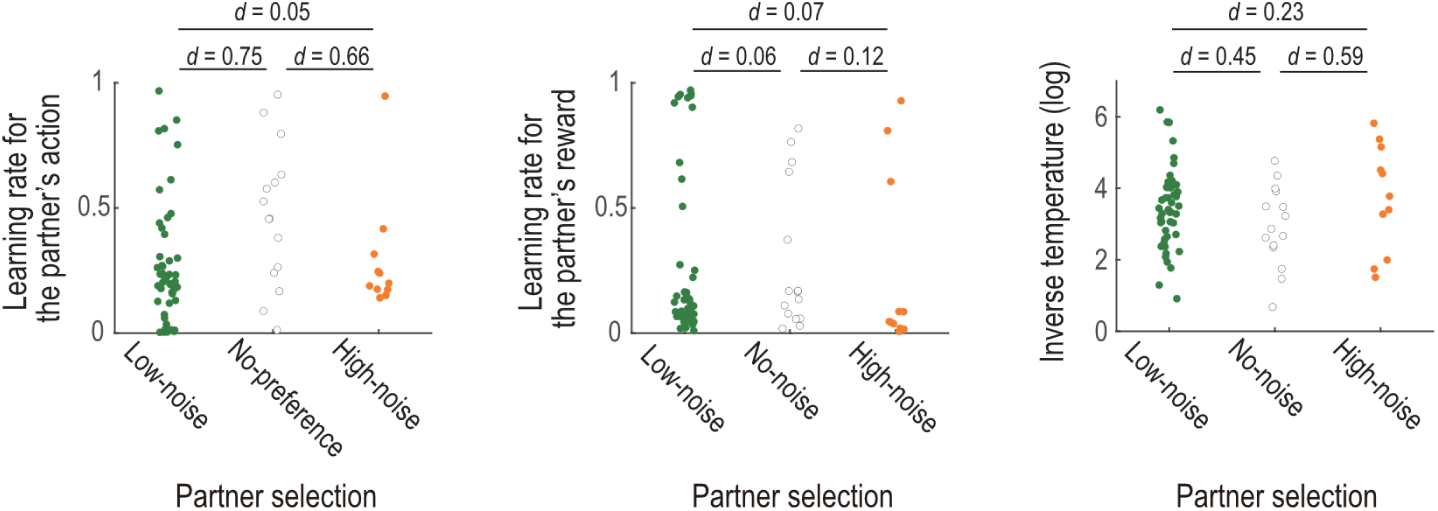
Estimated individual parameters from the full model: learning rate for the partner’s action (left), learning rate for the partner’s reward (middle), and inverse temperature (log scale; right), grouped by partner selection preference (Low-noise, No preference, High-noise). Effect sizes (Cohen’s d) are shown for pairwise group comparisons.

## Footnotes

1 In accordance with bioRxiv’s preprint policy, we replaced the original human face images with illustrative face drawings in Fig. 1. However, in the actual experiment, participants viewed real human faces sourced from a diverse, multi-racial dataset [51] to reduce potential biases related to sex and race. We plan to revert to using the original dataset images in the final published version.

